# Epigenetic heterogeneity shapes the transcriptional landscape of regional microglia

**DOI:** 10.1101/2024.08.08.607229

**Authors:** Alexander V. Margetts, Samara J. Vilca, Florence Bourgain-Guglielmetti, Luis M. Tuesta

## Abstract

Microglia, the innate immune cells in the central nervous system, exhibit distinct transcriptional profiles across brain regions that are important for facilitating their specialized function. There has been recent interest in identifying the epigenetic modifications associated with these distinct transcriptional profiles, as these may improve our understanding of the underlying mechanisms governing the functional specialization of microglia. One obstacle to achieving this goal is the large number of microglia required to obtain a genome-wide profile for a single histone modification. Given the cellular and regional heterogeneity of the brain, this would require pooling many samples which would impede biological applications that are limited by numbers of available animals. To overcome this obstacle, we have adapted a method of chromatin profiling known as Cleavage Under Targets and Tagmentation (CUT&Tag-Direct) to profile histone modifications associated with regional differences in gene expression throughout the brain reward system. Consistent with previous studies, we find that transcriptional profiles of microglia vary by brain region. However, here we report that these regional differences also exhibit transcriptional network signatures specific to each region. Additionally, we find that these region-dependent network signatures are associated with differential deposition of H3K27ac and H3K7me3, and while the H3K27me3 landscape is remarkably stable across brain regions, the H3K27ac landscape is most consistent with the anatomical location of microglia which explain their distinct transcriptional profiles. Altogether, these findings underscore the established role of H3K27me3 in cell fate determination and support the active role of H3K27ac in the dynamic regulation of microglial gene expression. In this study, we report a molecular and computational framework that can be applied to improve our understanding of the role of epigenetic regulation in microglia in both health and disease, using as few as 2,500 cells per histone mark.

**Figure 1.**
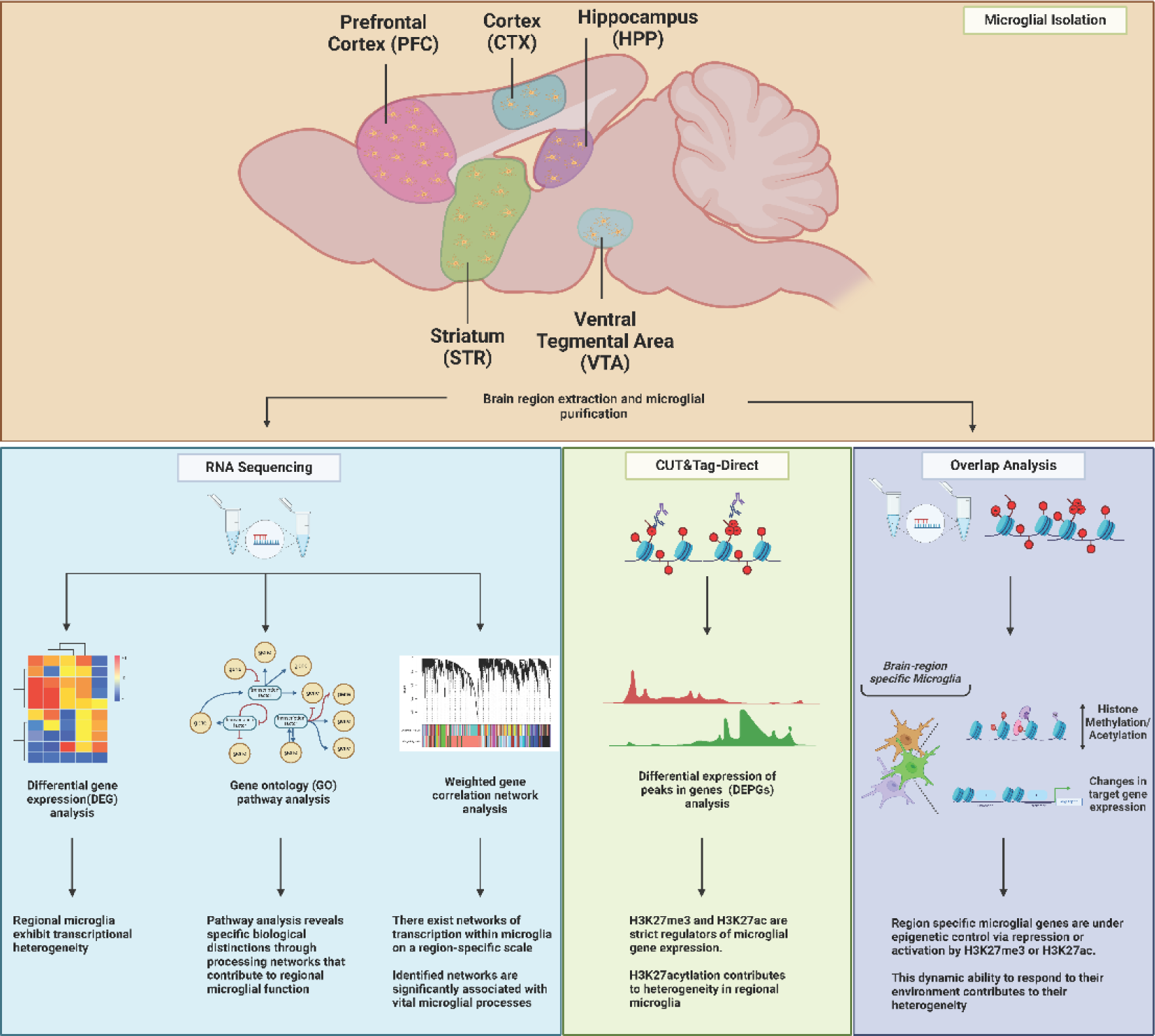
Pipeline of tissue processing and data analysis for the characterization of the microglial transcriptome and epigenome on a regional scale.

## Introduction

Microglia are the resident immune cells of the central nervous system (CNS). These cells contribute to various neurological functions throughout the lifespan, such as aiding in neurodevelopment by promoting neurogenesis, cellular differentiation, and shaping neuronal circuits [1, 2]. During adulthood, microglia participate in activity-dependent synaptic pruning, clearing cellular debris from dead and dying cells. Microglia continuously monitor the neural environment, responding to immunological insults or injury by sensing numerous pathogen and damage associated molecules. These reactive microglia are typically characterized by alternate morphologies and secretion of various cytokines (e.g., IL-6, IL-1b, and TNF-a) [1, 2]. Following this initial pro-inflammatory response, microglia then promote tissue repair and resolution of inflammation by releasing anti-inflammatory cytokines, chemokines, and neurotrophic factors [1, 2].

The brain is characterized by its rich cellular heterogeneity, as well as its complex circuitry that contributes to the functional specialization of each brain region. Indeed, microglial gene expression also demonstrates regional heterogeneity. For instance, hindbrain microglia exhibit higher expression of genes associated with clearance activity and immune signaling, while forebrain and midbrain microglia more robustly express genes related to synaptic plasticity and surveillance activity, although to varying levels [3–6]. Specifically, microglia marker genes *Cx3cr1* and *P2ry12* also display regional heterogeneity, showing higher expression in frontal regions compared to midbrain and hindbrain regions [7–9].

Interestingly, region-specific transcriptional differences in microglia may be influenced by epigenetic regulation through a suite of transcription factors, epigenetic modifying enzymes, and histone post-translational modifications (PTMs) [3, 10–12]. Indeed, studies suggest the homeostatic microglial transcriptome is regulated by master transcription factor PU.1 [13, 14] and chromatin modifications associated with active promoters (e.g., H3K4me3) and enhancers (e.g., H3K4me1/2 and H3K27ac) [15, 16]. Additionally, a recent study demonstrated that striatal and cerebellar microglial function are controlled by polycomb repressive complex 2 (PRC2) [11], which writes the repressive histone mark, H3K27me3. Altogether, these studies suggest microglia fine tune their gene expression to the surrounding neural environment through epigenetic modifications.

Indeed, the regions examined in this study are part of, and accessory to, the brain reward system and have been implicated in numerous psychiatric, neurodevelopmental, and neurodegenerative disorders [17–19]. Previous studies have suggested that microglia within these regions may contribute to disease through alteration of their transcriptional landscape [20–23]; however, epigenetic regulation of these cells is less known. Although much effort has been made to characterize the microglial epigenetic landscape in a brain region-specific manner, these efforts have been hampered by technical limitations. For instance, chromatin immunoprecipitation sequencing (ChIP-seq) and assay for transposase-accessible chromatin sequencing (ATAC-seq) have been first-line methods for epigenetic profiling across a broad range of cell types and tissues. In fact, both ATAC-seq and ChIP-seq have been invaluable in enabling the characterization of the microglial epigenome on a global scale [11, 24–31]. In this study we seek to build on these findings by better understanding how these effects can be measured on a regional scale. However, these methods require a significant amount of input (upwards of 500,000 cells) to obtain reliable results. Furthermore, while single-cell and single-nucleus sequencing have recently become the preferred methods to examine different cell populations, these are limited by their low sequencing depth. Therefore, a low-input method that retains a high signal-to-noise ratio with minimal sequencing depth (8-10M) is preferable to further examine the epigenetic landscape of brain region-specific microglia. Here, we utilize one such method, Cleavage Under Targets and Tagmentation (CUT&Tag-Direct), that can be used to profile the microglial epigenome from different brain regions with as few as 2,500 cells per histone mark. In this study, we examine the regional differences in naïve microglial gene expression across regions of the brain reward system and associate region-specific transcriptional profiles with repressive (H3K27me3) and permissive (H3K27ac) histone marks. In doing so, we describe a low-input molecular and computational framework, from isolation of primary microglia to preparation of RNA-seq/CUT&Tag-Direct libraries and computational analyses, that can be applied to study region-specific epigenetic and transcriptional changes in microglia both in health and disease models.

## Results

### Region-specific microglia are transcriptionally distinct

We first aimed to determine the transcriptional heterogeneity of microglia by brain region as previously observed [3, 11]. To this end, RNA from purified region-specific microglia were sequenced and differential gene expression analysis confirmed that microglia from the brain regions tested feature distinct transcriptional profiles. More specifically, we found that microglia from the hippocampus (HPP) are the most transcriptionally distinct when compared to somatosensory cortex (CTX), prefrontal cortex (PFC), ventral tegmental area (VTA) and the striatum (STR) (**Fig 2A**). In support, we found numerous differentially expressed genes between the brain regions examined (CTX, PFC, VTA, STR vs HPP: 2,681, 4,075, 4,020, 3,723, respectively, padj < 0.01) (**Fig 2B**, **Data S1**). Further, hierarchical clustering using the top differentially expressed genes across all regions shows region-specific transcriptional profiles (**Fig 2C**).

**Figure 2.**
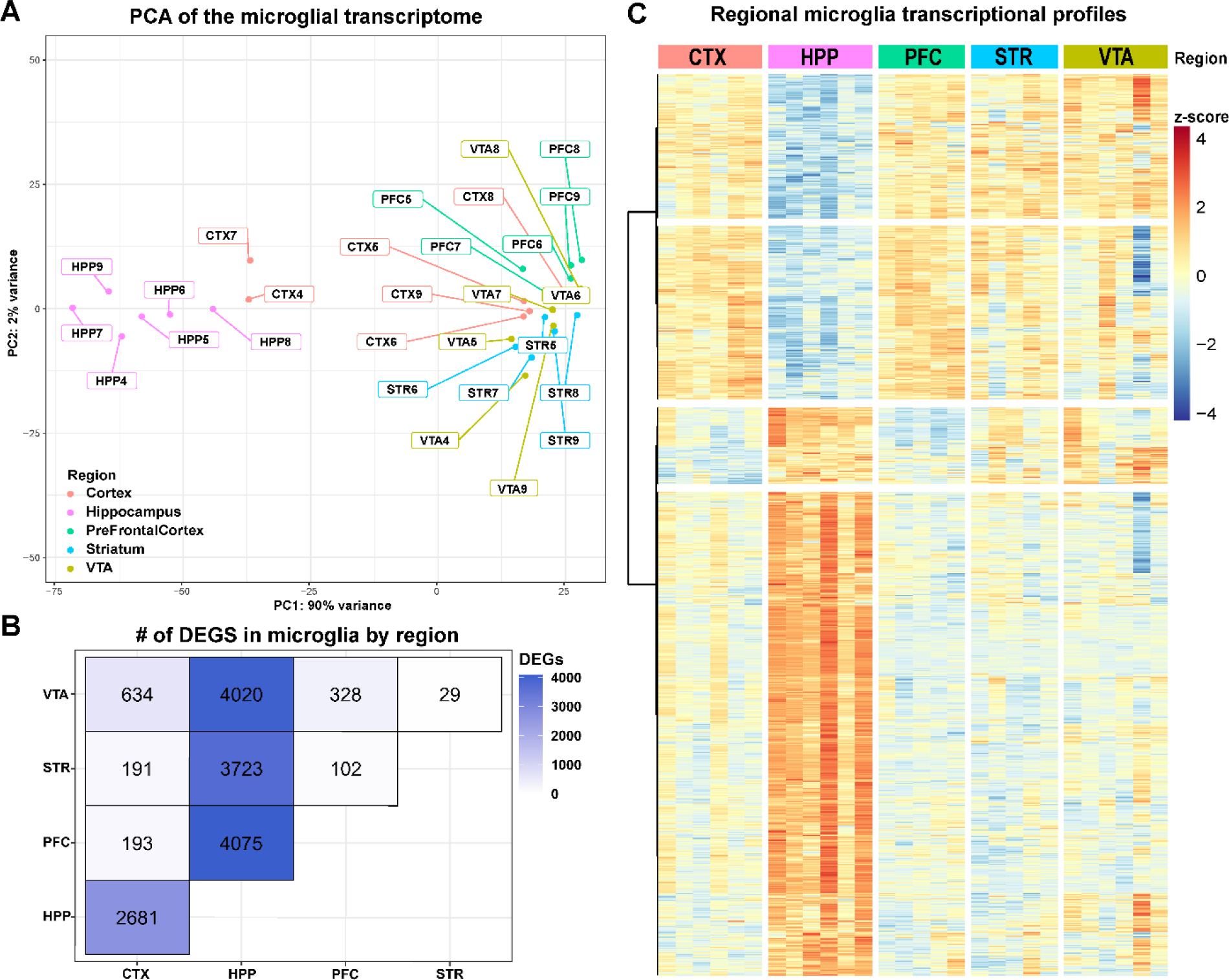
Microglia exhibit regional transcriptional heterogeneity. **(A)** Principal components analysis (PCA) indicates variation in baseline transcriptomics in hippocampal (HPP) microglia over other regions. Furthermore, we find **(B)** significant numbers of genes to be differentially expressed between each group in a pairwise comparison (n=5-6, padj <0.01). Of these significant genes we note **(C)** patterns of significant up- and down-regulation of genes in HPP microglia relative to other regions of interest.

### Pathway analysis reveals biological themes specific to regional microglia

To better understand the biological relevance of region-specific microglial transcriptomes, gene ontology (GO) pathway analysis was conducted on significantly differentially expressed genes. We found several significantly altered biological pathways which are specific to individual brain regions. Specifically, CTX microglia showed enrichment for pathways related to apical plasma membrane, external encapsulating structure organization, and extracellular structure organization when compared to VTA, as well as cilium processes when compared to STR (**Fig 3A**), suggesting that microglia in the somatosensory cortex may preferentially contribute to extracellular matrix (ECM) formation and overall cortical assembly and structure [32–34]. Additionally, we found microglia from the HPP were enriched for pathways such as axoneme when compared to the PFC, motile cilium when compared to the STR, and overall cilium organization when compared to the PFC, STR, and VTA (**Fig 3A**), indicating hippocampal microglia at baseline conditions may be highly surveilling and secretory compared to other brain regions [35]. Microglia from the PFC were significantly enriched for genes related to carbohydrate binding and leukocyte migration when compared to the HPP, MHC class II protein complex binding when compared to the STR, and humoral immune response when compared to both the STR and HPP (**Fig 3A**), consistent with other studies showing altered cytokine expression in microglia of the frontal cortex [36]. Furthermore, the locomotory behavior pathway was enriched in STR microglia when compared to the PFC (**Fig 3A**), supporting the importance of STR microglia in maintenance of dopaminergic circuitry [6, 37–39]. Additionally, the VTA was enriched for pathways related to chromosomal region, segregation, nuclear division, and regulation of cell cycle when compared to CTX, suggesting that microglia in the VTA may be more proliferative [6], whereas STR microglia were significantly enriched for pathways related to cytokine activity, regulation of cell-cell adhesion, and regulation of lymphocyte activation when compared to CTX microglia (**Fig 3A**). In addition, we find numerous genes that are expressed in a region-specific manner which also underscore these differences in regional microglia (**Fig 3B**).

**Figure 3.**
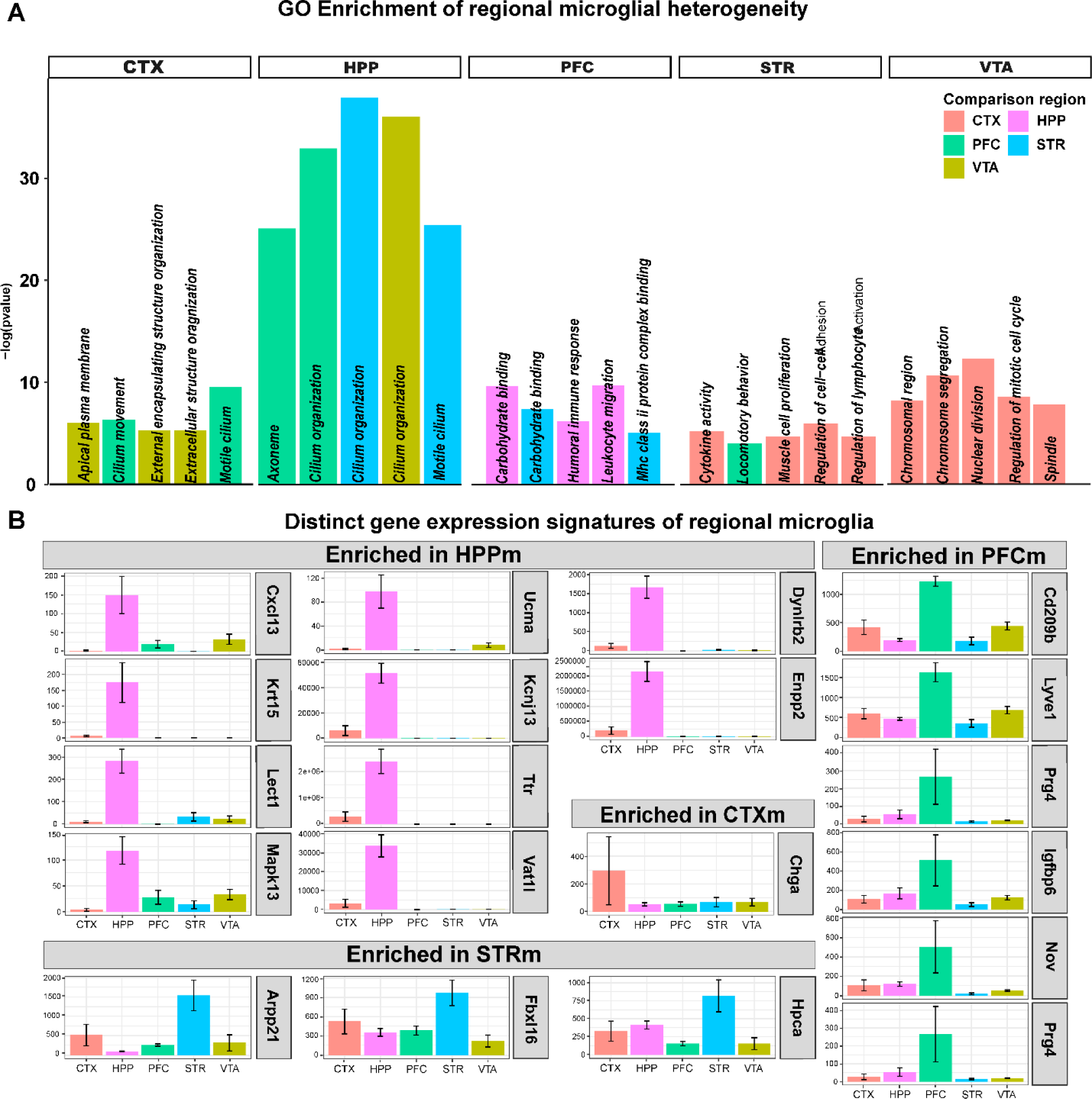
Transcriptional heterogeneity is linked to region specific gene expression and underlying functional differences. **(A)** Significantly enriched Gene Ontology (GO) pathways for each region studied (n = 5-7, padj_genes_ < 0.05; padj_pathways_ < 0.01) in a pairwise fashion. **(B)** Normalized gene expression plots highlight significantly enriched genes that are highly expressed in each region (n = 5-7, padj_genes_ < 0.05, Error bars represent SEM).

### Weighted gene-network correlation analyses reveal region-specific networks of microglia gene expression

Following differential gene expression analyses, we sought to better characterize the regional differences observed in microglia by identifying modules of expression. To this end, a weighted gene correlation network analysis was conducted and identified 8 modules which show significant mean differential expression across brain regions (**Data S2**). Of these 8 modules, the “salmon” (ME1), “blue” (ME4), “plum” (ME5) and “darkolivegreen” (ME19) modules were selected for further analysis, as they were the most differentially expressed (**Fig 4A**). To understand the biological relevance of these genes, we first filtered the modules for genes with high module membership based on *p*-value correlations and identified hub genes central to each module (**Fig 4C-F**). GO pathway analyses on highly modular genes, those representing the top 1% of modular membership, revealed several differentially expressed pathways between regions (**Fig 4B**). Interestingly, we found that ME1, which is mainly involved in movement and cellular structure, is significantly enriched in hippocampal microglia relative to other regions, while ME5 was significantly depleted in HPP microglia, indicating that GTPase enzymatic activity may be higher in microglia from other regions compared to HPP. Additionally, STR and VTA microglia show enrichment of pathways associated with an immune-activated phenotype including responses to bacteria and stimuli, identified in ME4. Finally, ME19 was slightly enriched in HPP, STR and VTA microglia with higher representation of genes related to cell-cycle phase transition.

**Figure 4.**
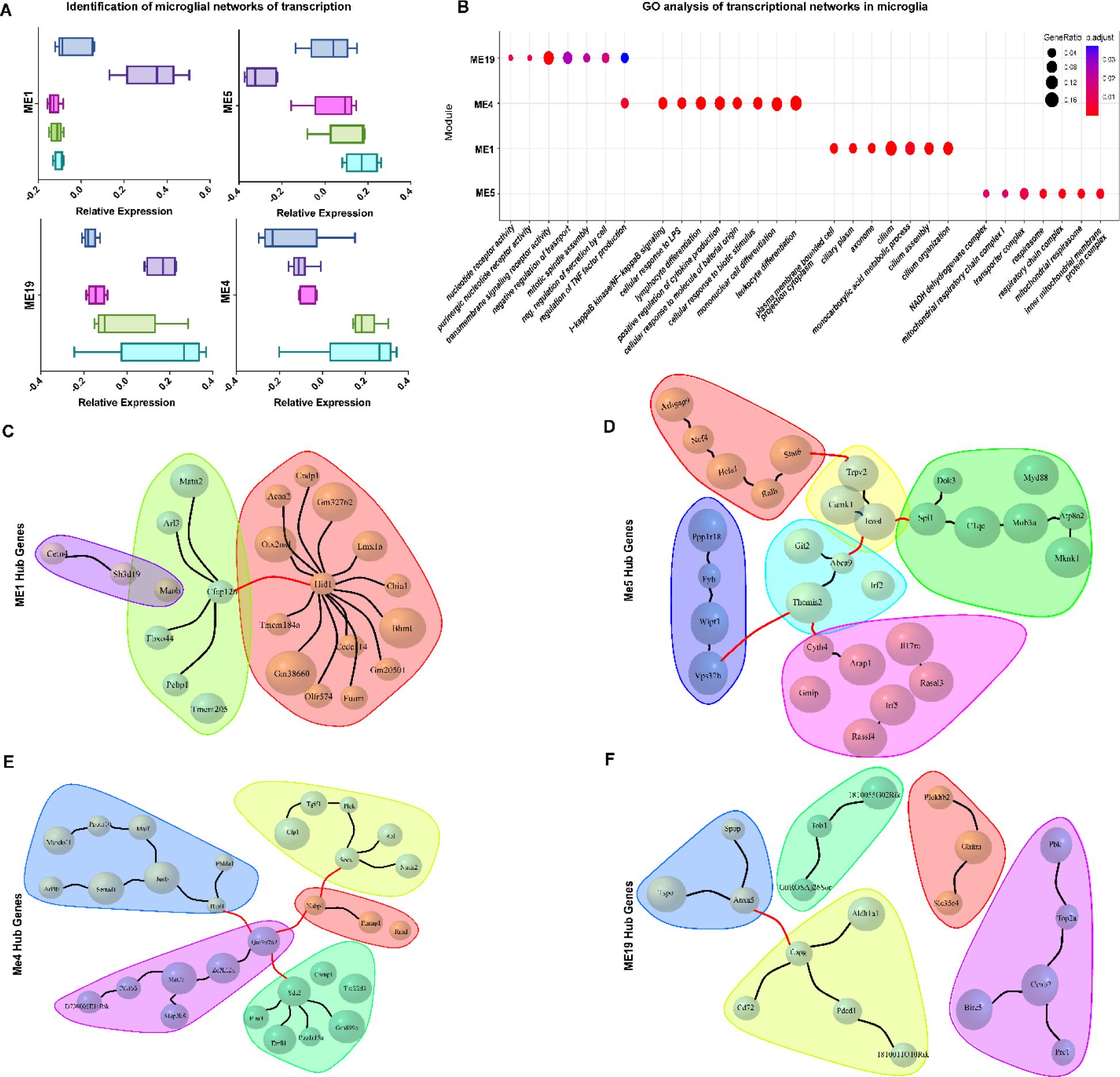
WGCNA analysis reveals networks of transcription in regional microglia. **(A)** Significantly differentially expressed modules identified by weighted gene correlation network analysis (limma, ME1: padj < 0.00001; ME5: padj < 0.001; ME19: padj < 0.01; ME4: padj < 0.01) **(B)** GO pathways enriched in highly modular genes for each of the significantly enriched modules (padj < 0.05). Identified hub genes from high modular membership genes (top 1% of genes in each module with module membership *p-*value < 0.05) in the **(C)** ME1 module, **(D)** ME5 Module, **(E)** ME4 Module, **(F)** ME19 Module. (Colors indicate hub genes with closest relationships. Red lines indicate genes with significant connections (Pearson correlation, *p*-value < 0.05))

Taken together, these data support the hypothesis that regional neural microenvironments may regulate microglial gene expression, and that these transcriptional variations enable microglia to execute their unique homeostatic functions across different brain regions. While we identified hippocampal microglia as most transcriptionally distinct from other region-specific microglia, the epigenetic factors that contribute to such heterogeneity remain unclear.

### CUT&Tag-Direct enables histone profiling on minimal primary microglial samples

H3K27me3 is a histone PTM associated with repression and is often found at the transcriptional start site (TSS) and throughout the gene bodies of repressed genes [40, 41]. Conversely, H3K27ac is associated with active transcription and is mainly deposited at, or near the TSS or promoter regions of expressed genes [42]. In order to elucidate the epigenetic regulation of region-specific microglial genes, we conducted CUT&Tag-Direct [43] for H3K27me3 and H3K27ac. Due to the limited number of microglia obtained from tissue extractions, we used CUT&Tag-Direct, a modification of the original protocol for limited cell numbers. To ensure accurate epigenetic mapping, an equal number of designer nucleosomes (dNucs) against a range of histone methylation marks were spiked into each reaction. When compared to other methylation dNucs, the H3K27me3 antibody used for these assays showed high specificity (< 10% non-specific binding) (**Fig S1A**). Indeed, immunohistochemical analysis revealed that the antibodies used were specific and that microglia contain sufficient nuclear material regulated by these marks (**Fig 5A**). Samples yielded robust alignment to the mm10 genome (87% - 97% aligned reads) (**Data S3**). After merging technical replicates, which showed high levels of similarity (**Fig S1B**), peaks were called using SEACR[44], which identified upwards of 150,000 significant peaks in each region for both H3K27me3 and H3K27ac (**Data S3**).

**Figure 5.**
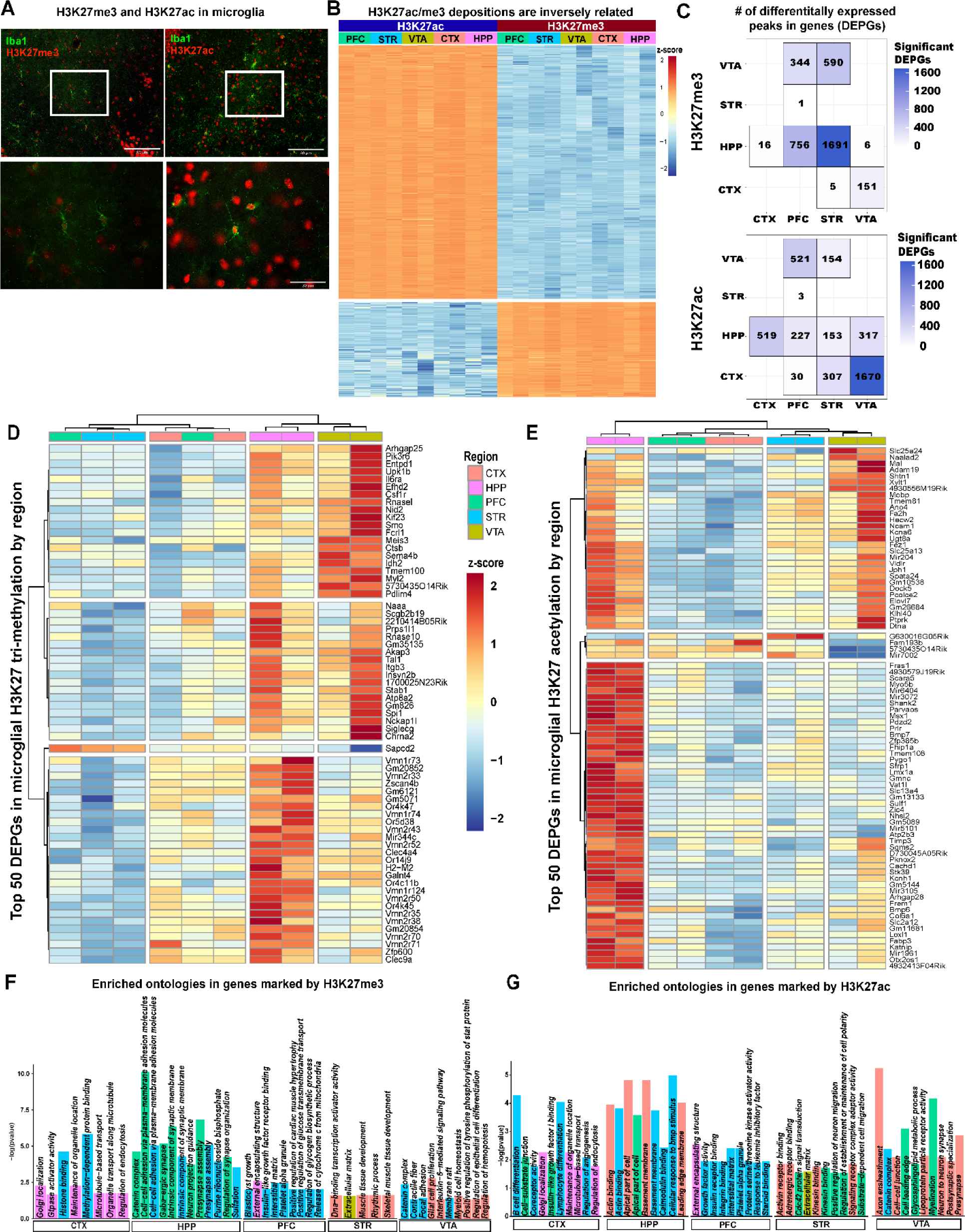
Differential peak deposition analysis. **(A)** Representative immunohistochemical images of mouse HPP microglia, using the microglial marker Iba1 and H3K27me3 (left) and H3K27ac (right) antibodies that were used in CUT&Tag-Direct. **(B)** Representative heatmap of fragments in the TSS or promoter (H3K27ac) or TSS, promoter or exons (H3K27me3) of genes with significantly called peaks (SEACR). Number of significantly differentially deposited peaks in genes for **(C)** H3K27me3 and H3K27ac in a region to region pairwise comparison. Heatmaps of the top differentially deposited peaks in genes for **(D)** H3K27ac and **(E)** H3K27ac (−1.25 < LFC > 1.25, padj <0.05, LfcSE <1). GO pathway analysis results for **(F)** H3K27me3 peaks enriched in genes and **(G)** H3K27ac peaks enriched in genes.

### Epigenetic contributions to microglial heterogeneity

To determine the contribution of these histone PTMs to transcriptional differences in region-specific microglia, we generated fragment counts within genes related to the histone marks of interest and conducted differential peak deposition analysis using DESeq2. In agreement with previous studies[45], H3K27ac and H3K27me3 are not often simultaneously present at gene promoters (**Fig 5B**). Interestingly, H3K27ac appears to vary by region, suggesting this mark may regulate expression of many genes, whereas H3K27me3 shows a more stable deposition pattern (**Fig S2A**). H3K27me3 deposition varied greatly in HPP microglia over other regions, and a majority of significantly differential depositions in genes (DEPGs) appeared in HPP versus STR (1,691 genes containing differential depositions, padj <0.05). Additionally, HPP microglia greatly differ when compared to PFC microglia (756 genes containing differential deposition, padj <0.05). Furthermore, PFC and STR microglia exhibit differential deposition of H3K27me3 when compared to VTA (344, 590 and 151 genes, padj < 0.05) (**Fig 5C, Data S3**). These patterns of H3K27me3 deposition in microglia suggest epigenetic heterogeneity in each region (**Fig 5D**). GO pathway analysis revealed that H3K27me3 deposition is significantly different in genes related to synaptic processes including pre-synapse assembly, organization and structure, as well as GABA receptor complexes when comparing HPP to both STR and PFC microglia (**Fig 5F, Data S3**). Microglia from the VTA exhibit significant differential deposition of H3K27me3 in pathways related to extracellular matrix and basement membrane (vs PFC), actin-filament and methylated histone or protein binding (vs HPP), glial cell proliferation, histone deacetylase binding and multiple differentiation pathways (vs CTX), and actin cytoskeleton (vs STR) (**Fig 5F, Data S3**).

Unlike H3K27me3, H3K27ac is a transient mark of active gene transcription [42, 46]. Indeed, significant differential deposition of H3K27ac is far more abundant than that of H3K27me3 (**Fig 5C, Data S3**). Similar to H3K27me3 depositions, H3K27ac varied greatly in the HPP over other regions. H3K27ac in hippocampal microglia varies greatly when compared to microglia from the CTX, PFC, STR, and VTA (519, 227, 153, 317 genes containing differential deposition, respectively, padj < 0.05). However, the majority of significantly DEPGs are found in the VTA as compared to the CTX (1,670 genes containing differential depositions, padj <0.05). Furthermore, when compared to the VTA, microglia from the PFC and STR exhibit differential regulation of 521 and 154 genes by H3K27ac, respectively (padj <0.05). Mirroring H3K27me3, the deposition pattern of H3K27ac in microglia indicates epigenetic regulation of the transcriptional heterogeneity observed across regions (**Fig 5E, Data S3**). GO pathway analysis highlights this heterogeneity, as HPP microglial H3K27ac deposition is significantly enriched in genes involved in actin filament and binding as well as extracellular matrix (vs CTX), cell to cell signaling and glial cell migration (vs STR), cell projection and apical plasma membrane (vs VTA), and actin filament related processes (vs PFC) (**Fig 5G, Data S3**). Differential H3K27ac deposition in the VTA compared to the CTX and PFC indicates significant enrichment in genes related to myelination, as well as axon and neuron ensheathment (**Fig 5G, Data S3**).

### Correlating regional microglia gene expression with abundant gene expression

To better understand the overall transcriptional regulation of microglial genes by H3K27me3 and H3K27ac, we conducted a gene-overlap analysis. Here, we sub-set gene expression into lowly expressed (FPKM < 1), moderately expressed (FPKM > 1, FPKM <10) and highly expressed (FPKM > 10) and overlapped these with significantly called peaks found at TSS (for both H3K27me3 and H3K27ac) or exonic regions (for H3K27me3) in protein coding genes. By doing so, we found significant overlap of H3K27ac (active transcription) in all regions with both moderately expressed and highly expressed genes. On the other hand, we found significant overlap of H3K27me3 (repression) in all regions with lowly expressed genes (**Fig S2A**). Furthermore, 1960, 1190, 1552, 1609 and 687 lowly expressed genes are significantly correlated with H3K27me3 deposition, while 738, 667, 897, 739, and 749 highly expressed genes are significantly correlated with H3K27ac deposition in the PFC, STR, VTA, HPP and CTX, respectively (**Fig S2B**).

To identify the genes contributing to regional heterogeneity in microglia, the gene lists were then filtered to include only genes unique to each region and containing deposition of each histone modification. Remarkably, 151, 117, 267, 173 and 162 highly expressed genes and 226, 215, 270, 279, and 202 moderately expressed genes overlap with significantly called peaks for H3K27ac that are unique to the PFC, STR, VTA, HPP and CTX, respectively. Additionally, we found 229, 146, 213, 186 and 194 lowly expressed genes overlap with significantly called peaks for H3K27me3 that are unique to the PFC, STR, VTA, HPP and CTX, respectively (**Fig S2B**). Indeed, while peaks may be present in the promoter-TSS or gene bodies of these region, thereby indicating uniqueness, these results indicate regional specificity of microglial gene expression that is highly correlated with histone PTMs, with the most genes being regulated by H3K27ac in the VTA (**Fig S2B**). While these patterns suggest epigenetic regulation of genes in a region-specific manner, finding the biological relevance of such regulation requires further characterization.

### Region-specific epigenetic regulation of microglial genes

We next sought to obtain an overarching understanding of gene regulation via H3K27me3 and H3K7ac in microglia based on neural environments by overlapping DEGs that contain significant deposition of specific histone PTMs. We find a number of significant overlaps when comparing regions (**Fig 6A, C**). Indeed, in agreement with our transcriptomic data, most significant overlaps are found when comparing HPP microglia to other regions. GO analysis of overlapping genes indicates that epigenetic control of genes by H3K27ac in hippocampal microglia are enriched for pathways related to actin regulatory processes, BMP signaling pathway, and kinases (**Fig 6B**). Hippocampal microglia also show significantly downregulated gene expression (due to high H3K27me3 deposition) related to various immune, proliferation and differentiation pathways (**Fig 6B**). These findings highlight the need for this type of analysis, as merging epigenetic and transcriptomic data underscores differences in microglia across the brain.

**Figure 6.**
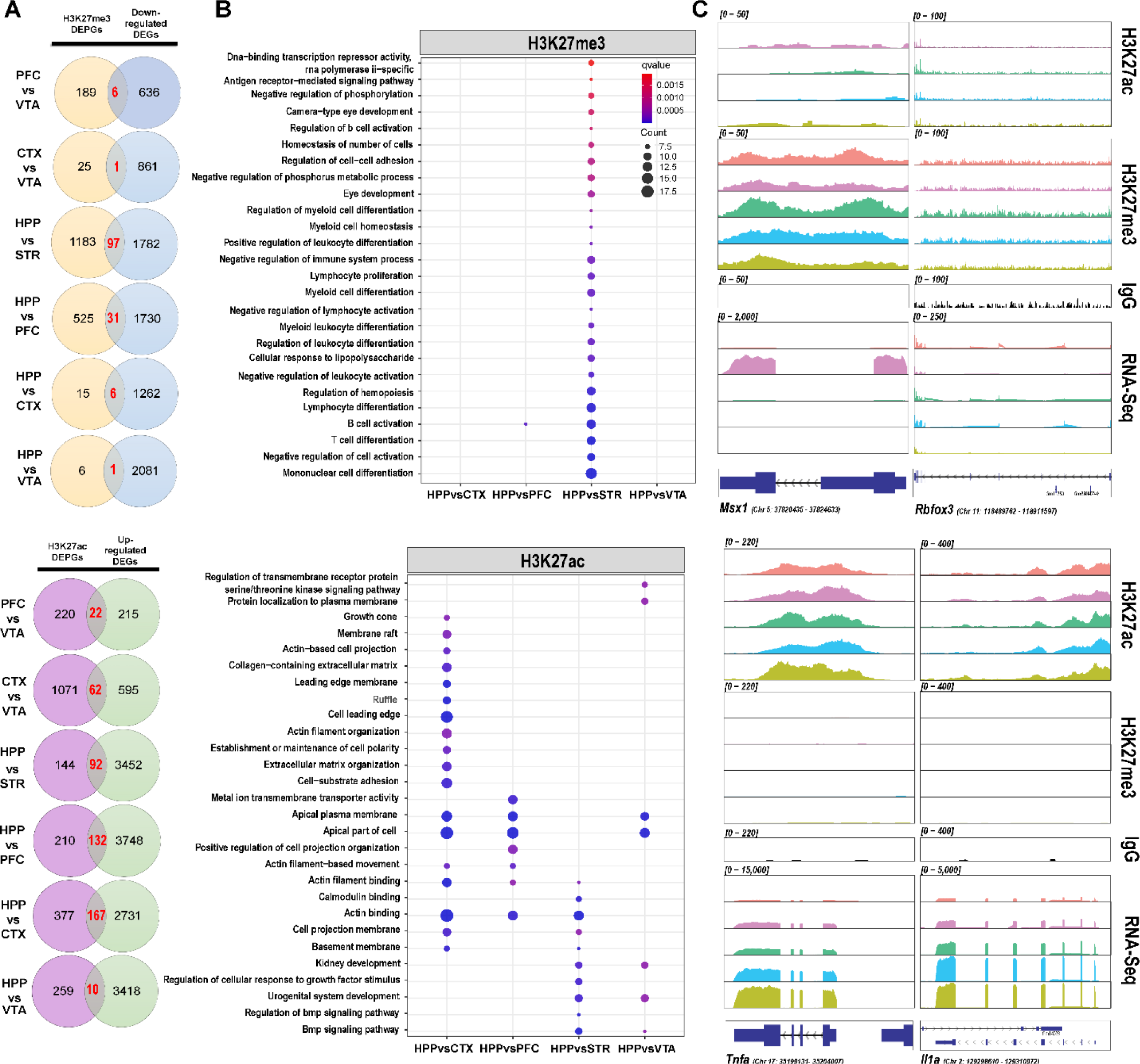
Differential peak deposition analysis. **(A)** Venn diagrams showing the number of significant H3K27me3 depositions in genes (DEPGs) (n = 2, padj < 0.05) with significant upregulated genes (DEGs) (n = 2, padj < 0.05) for each regional comparison. (Red numbers indicate overlapping DEPGs and DEGs for each comparison) **(B)** GO analysis dot plots utilizing overlapping genes as input (padj < 0.10) **(C)** Representative IGV plots of CUT&Tag-Direct and RNA-sequencing normalized BigWigs.

## Discussion

Microglia are the primary immune cells of the CNS and play a key role in development and homeostasis, as well as disease states [47]. Microglial activity largely depends on communication with neurons and other glia, which can induce both transcriptional and epigenetic changes in microglia. While previous research has examined whole-brain changes in microglial transcriptional profiles, recent studies have interrogated these cells in a region-specific manner. A great focus has been placed on the various transcriptional responses of microglia to different stimuli, yet homeostatic transcriptional differences across the brain have only recently been investigated [3]. Furthermore, epigenetic regulation, which can drive transcriptional heterogeneity, has yet to be fully explored in microglia. This may be due, in part, to the inability to obtain sufficient material for both bulk RNA-seq and epigenetic assays from genetically defined cell populations within a single brain region. Our experimental approach, which optimizes CUT&Tag-Direct in conjunction with low-input RNA-seq, enabled us to profile region-specific primary mouse microglia without the need for large numbers of cells or single-cell sequencing techniques that can be burdened by shallow coverages.

Here, we explored the epigenetic regulation of transcriptional heterogeneity in microglia across five brain reward regions in mice (somatosensory cortex, prefrontal cortex, striatum, hippocampus, and ventral tegmental area). This is the first study of its kind to generate both transcriptional and epigenetic data from primary microglia from the same animals. In this study, we found robust gene expression differences in microglia across all five mouse brain regions queried, highlighting their transcriptional diversity (**Fig 2A-C**). We also identified several regionally expressed genes which may contribute to functional differences in microglia (**Fig 3B**). In our data, GO analysis indicated that naïve hippocampal microglia at baseline are enriched for the aforementioned pathways as compared to PFC, STR and VTA microglia (**Fig 3A**), implying microglia in the hippocampus may form cilia to increase their sensing ability and secretory function [35]. Indeed, previous studies have shown that decreased expression of genes related to axoneme assembly and cilium organization pathways in the hippocampus is associated with Alzheimer’s Disease and decreased ramifications and impaired sensing in microglia [35, 48]. We also observed that microglia from the VTA express genes reflective of increased proliferation when compared to CTX microglia, which is consistent with previous studies [6]. Overall, profiling transcriptional homeostatic differences in naive microglia will be key to understanding regional variability under experimental or disease conditions.

Furthermore, in order to identify networks of transcription, or gene co-expression, which cannot be identified using DEG analysis, we conducted WGCNA. WGCNA revealed the existence of subsets of genes that share high expression in some regions, while being lowly expressed in others. Importantly, this study identifies four modules of interest that expand our knowledge of microglial transcriptional heterogeneity. Of note, we again find a number of axoneme, and cilium related pathways enriched in the hippocampus versus other regions. We also found that mitosis and numerous immune related pathways are enriched in microglia from the STR and VTA (**Fig 4A-B**), which aligns with previous studies indicating that VTA microglia show a unique reactive phenotype and may be more susceptible to age-related dysfunction [6, 49, 50], as well as other studies that show higher expression of homeostatic and immune surveillance genes in striatal microglia [11]. From this analysis, we also find that microglia from the CTX and PFC share similar expression across almost all modules, generally expressing genes related to transmembrane signaling receptor activity, indicating a more homogeneous population of microglia in these two regions (**Fig 4A**), also consistent with previous studies [3, 6]. While these transcriptional differences have been previously reported, this study sought to further associate these changes with epigenetic modifications.

Previously, ChIP-seq and ATAC-seq have been used to characterize microglia on a global scale [10, 11, 30, 31], while more recent studies have conducted single-cell RNA-seq and single-cell ATAC-seq to analyze gene expression and chromatin accessibility [51–57]. Unfortunately, the input required to conduct ChIP-sequencing is often insufficient for regional analysis without requiring prohibitive numbers of animals. While ATAC-seq can identify open (active) regions of chromatin, specific histone PTMs cannot be measured. This study establishes a viable methodology for conducting epigenetic profiling on region-specific microglia, a cell population of very limited yield. Indeed, CUT&Tag-Direct yielded robust, high signal-to-noise data and enabled the identification of upwards of 250,000 histone peaks in some samples (**Data S3**), along with high mapping rates and specificity for the marks studied (**Fig S1**). By focusing on two histone PTMs known to regulate gene expression, H3K27ac and H3K27me3, this study identified uniquely regulated genes across specific brain regions in mice (**Fig 5B**). Furthermore, as previous studies have shown, we found that H3K27ac and H3K27me3 do not occupy the same promoters and genes simultaneously, due to their opposing effects on gene expression (**Fig 4A**) [45].

This study also uncovered that while histone mark deposition varies by region, these align with gene expression (**Fig 5A**). Most of the transcriptional variation in our dataset was due to regional differences in H3K27ac deposition (**Fig S3A**). By conducting RNA-sequencing in conjunction with CUT&Tag-Direct, this study identified region-specific gene expression that is significantly associated with deposition of either H3K27me3 or H3K27ac. Furthermore, by identifying genes that contained significant depositions of H3K27ac or H3K27me3, we were able to correlate this with regional expression of genes (**Fig 5A**). In fact, many genes identified in our WGCNA analysis were found to have reasonable deposition of each mark indicating there may be regulation of these regional transcriptional networks (**Fig S3B**). These overlapping features indicated that the presence of these marks may directly impact gene transcription in a region-specific manner. Indeed, we not only uncovered that *Cx3cr1* and *Tnf-a* contain depositions of these marks, indicating regulation, but we found unique genes specific to each region (**Fig 5B**). Given the known role of H3K27me3 in cell fate determination, we were not surprised that there were far fewer H3K27me3-repressed genes that were unique to each region [58]. On the other hand, we found between 17% and 30% of genes that were unique and highly expressed in each region are associated with H3K27ac expression (**Fig 5B**). Taken together, these results suggest that microglial transcriptional heterogeneity is under regulation, at least in part, by deposition of repressive and permissive histone marks in H3K27.

This study furthers our understanding of epigenetic regulation of regional microglial gene expression by identifying genes containing significant depositions of H3K27ac or H3K27me3, and which were significantly upregulated or significantly downregulated, respectively (**Fig 6A**). Notably, these differences identify hippocampal microglia as being the most heterogeneous relative to the other regions of the brain reward system that we examined. Interestingly, when GO analysis was conducted on these genes, we found that H3K27ac in hippocampal microglia drove expression of genes involved in cellular transport and cellular structure, while increased presence of H3K27me3 in downregulated genes was identified in immune response-related pathways such as B-cell activation and responses to stimuli such as LPS (**Fig 6B**). Further, we note that concomitant decreases in H3K27me3 and increases in H3K27ac can drive differential gene expression of certain genes, such as *Msx1*, across regions (**Fig 6C**). These data suggest that hippocampal microglia may be more homeostatic, with broad deposition of H3K27me3 in genes related to immune pathways, while deposition of H3K27ac occurs in genes necessary for surveillance when compared to other regions of the brain reward system.

Sex is an important consideration when studying microglia given that neuroimmune system development is regulated by this biological variable [59, 60] and microglial transcriptional profiles have been shown to differ in males and females [3]. Indeed, while we focused on two key post-translational modifications, namely H3K27ac and H3K27me3, there are numerous other epigenetic contributions to transcriptional control in microglia [12]. Further studies focusing on sex differences within and beyond the regions studied here, including additional post-translational histone modifications, will be necessary to more fully understand the contributions of epigenetic diversity to transcriptional regulation of microglial regional heterogeneity.

Taken together, these data combine transcriptional and epigenetic datasets from region-specific primary mouse microglia and highlight the molecular heterogeneity that exists in these cells across the brain reward system. Most importantly, this study establishes a methodology for studying the epigenetic and transcriptional changes that occur in cell populations of limited yield. Such populations may be limited by their anatomical location, or by their limited numerical representation relative to functional importance (i.e. dopamine neurons). Furthermore, application of this molecular and computational framework may also be useful for understanding the epigenetic mechanisms that undergird phase, or time-dependent changes in gene expression associated with various psychiatric, neurodegenerative or neoplastic diseases.

## Materials and Methods

### 1. Animals

Male C57BL/6J mice (12–16-week-old, ∼25-30 g; Jackson Laboratories, Bar Harbor, ME; SN: 000664) were housed in the animal facilities at the University of Miami Miller School of Medicine. Mice were maintained on a 12:12 h light/dark cycle and were housed three to five per cage. Animals were provided with food and water *ad-libitum*. All animals were maintained according to National Institutes of Health (NIH) guidelines and Association for Assessment and Accreditation of Laboratory Animal Care (AAALAC) accredited facilities. All experimental protocols were approved by the Institutional Animal Care and Use Committee (IACUC) at the University of Miami Miller School of Medicine.

### 2. Microglial Isolation

Mice were anesthetized with isoflurane and perfused through the ascending aorta with 1X phosphate buffer saline (PBS; pH 7.4; ThermoFisher, 10010023) plus heparin (7,500 USP units). Brain regions from 4 mice (**Fig 1**) were dissected and combined, then transported on ice in Hibernate A Medium (Gibco, A1247501). Tissue was then dissociated on the gentleMACS Octo Dissociator (Miltenyi Biotec, #130-096-427) using the Adult Brain Dissociation Kit (Miltenyi Biotec, #130-107-677) according to manufacturer’s instructions. All steps after initial dissociation were performed on ice and all tubes were prechilled. The resulting single cell suspension was incubated with anti-mouse CD11b (microglia-specific) magnetic MicroBeads (Miltenyi Biotec, #130–093–634) and microglia were positively selected via column purification (Miltentyi Biotec, #130-042-201). The eluted fraction containing microglia was then centrifuged for 3 min at 600 *x g* and resuspended in 110 µL of PBS (ThermoFisher, 10010023). 10 µL was reserved for cell counting on the EVE Countess Automated Cell Counter (NanoEntek), and the remaining 100 µL was split: 30 µL for RNA extraction, 70 µL for CUT&Tag-Direct.

### 3. Brain Perfusion and Fixation

Mice were anesthetized with isoflurane and perfused through the ascending aorta with 1X phosphate buffer saline (PBS; pH 7.4; ThermoFisher, 10010023) plus heparin (7,500 USP units) (DVR University of Miami), followed by fixation with 4% paraformaldehyde (PFA; Sigma Aldrich, 1003543951) in PBS. Brains were collected, postfixed overnight in 4% PFA, and then stored in 30% sucrose with 0.05% sodium azide (Sigma Aldrich, 1003124924) in PBS. All brains were cut into 35 µm coronal sections on a cryostat, and floating sections were stored in PBS with 0.02% sodium azide at 4°C until processing for immunohistochemistry.

### 4. Fluorescence Immunolabeling

Floating sections were processed for fluorescent immunostaining of microglia. Sections were rinsed in PBS and then blocked for 1 hour at room temperature in Blocking Buffer consisting of 10% normal donkey serum (Jackson ImmunoResearch, 017-000-121), 0.5% Triton X-100 (Sigma Aldrich, T8787), and PBS. Thereafter, sections were incubated in primary antibody in Blocking Buffer at 4°C overnight. The primary antibodies used were as follows: goat anti-Iba1 (1:500, NB1001028, Novus Biologicals), rabbit anti-H3K27me3 (1:500, 9733S, Cell Signaling Technology), and rabbit anti-H3K27ac (1:160, 8173, Cell Signaling Technology). On day 2, the sections were rinsed 3 times with PBS and incubated with the following secondary antibodies: Alexa 488 donkey anti-goat (1:500, A32814, Invitrogen) and Alexa 568 donkey anti-rabbit (1:500, A10037, Invitrogen). Sections were incubated with secondary antibodies in PBS with 2% normal donkey serum for 2 hours at room temperature in the dark. Next, sections were rinsed 3 times with PBS, mounted on slides with ProLong Diamond Antifade Mountant with DAPI (Invitrogen, P36962) and cover slipped. Images were acquired using the VS120 Olympus slide scanner housed in the University of Miami Miller School of Medicine’s Analytical Imaging Core Facility (AICF). Controls included processing the secondary antibodies alone to verify background staining, processing each primary with the secondary antibody to verify laser-specific excitation, checking for autofluorescence in an alternative laser channel with tissue lacking that laser-specific probe, and using sequential scanning. Fluorescent images were viewed in OlyVIA (Olympus, Ver.2.9.1), where only brightness and/or contrast levels were adjusted after acquisition were imposed across the entire image. All antibodies used have been previously validated for the intended applications, as per manufacturer. For all representative images of qualitative data, the immunolabeling experiment was successfully repeated in 3 animals.

### 5. RNA-seq

30 µL of isolated microglia was added to 350 µL RLT plus buffer (Qiagen, 1053393) for extraction and purification of total RNA according to manufacturer’s instructions using the Qiagen AllPrep DNA/RNA Mini Kit (Qiagen, 80204). Total RNA input was normalized and NGS libraries were prepared using NEBNext Single Cell/Low Input RNA Library Prep Kit for Illumina (New England BioLabs, E6420S) according to the manufacturer’s instructions. Sequencing was performed on an Illumina NovaSeq6000 platform (150×150bp PE) targeting 30 million reads per sample by the University of Miami Center John P. Hussman Institute for Human Genomics sequencing core facility.

### 6. RNA-seq Analysis

All RNA-seq data used in this study were mapped to the mm10 genome. Prior to mapping, raw RNA-seq datasets were first trimmed using TrimGalore (v.0.6.7) [61] with cutadapt (v.1.18) [62]. Illumina sequence adaptors were removed, the leading and tailing low-quality base-pairs (fewer than 3) were trimmed. Next, reads with a length of at least 20-bp were mapped to the genome using STAR (v.2.7.10a) [63] with the following parameters: –outSAMtype BAM SortedByCoordinate –outSAMunmapped Within –outFilterType BySJout –outSAMattributes NH HI AS NM MD XS –outFilterMultimapNmax 20 –outFilterMismatchNoverLmax 0.3 --quantMode TranscriptomeSAM GeneCounts. The resulting bam files were then passed to StringTie (v.2.1.7) [64] to assemble sequenced alignments into estimated transcript and gene count abundance given the Gencode GRCm38 (NCBI) transcriptome assembly.

#### 6.1 Differential Gene Expression Analysis

The R/Bioconductor DESeq2 (v.1.34.0) [65] package was used to detect the differentially expressed genes between region-specific microglia. Only genes with transcript abundance > 50 across samples, and adjusted *p*-value (padj) < 0.05 were considered as differentially expressed.

#### 6.2 Functional Enrichment Analysis

The R/Bioconductor clusterProfiler (v.4.2.2) [66, 67] package was used to perform Gene Ontology (GO) analysis. Only the GO terms and pathways with a padj < 0.05 following false discovery rate (FDR) correction were considered, with focus given to those showing positive enrichment only (i.e., log_2_foldchange (log_2_FC) > 1). The R/Bioconductor package rrvgo (v1.6.0) [68] was utilized to reduce GO terms to parent terms for clarity. The associated GO and pathway enrichment plots were generated using the ggplot2 package (v.3.4.2). Heatmaps were generated using the R/Bioconductor package pheatmap (v.1.0.12). All the other plots were generated using the ggplot2 package.

#### 6.3 Weighted Gene Co-Expression Analysis (WGCNA)

The R package WGCNA [69] was used to conduct gene co-expression analysis. Briefly, count matrices were loaded into R and normalized using a variance stabilized transformation (VST). Samples were checked for goodness and manual network construction using a bicorrelation, soft thresholding power of 7, a signed-hybrid network and a signed TOM was conducted. The minimum module size was set at 30 genes and modules were merged based on similarity and assigned a color name. The limma package (v.3.50.3) [70] was used to conduct statistical analysis to identify which modules were significantly differentially expressed (padj > 0.05) by brain region based on average expression of all genes in that module following FDR multiple testing correction. Module expression was plotted with ggplot2 (v.3.4.2). Module membership was calculated and the top 1% of most modular genes were selected and GO pathway enrichment was conducted on them using the R/Bioconductor package clusterProfiler (v.4.2.2) [66, 67] against a background containing all genes mapped to relevant modules with an FDR multiple testing correction. Heatmaps were generated using the R/Bioconductor package pheatmap (v.1.0.12) by generating lists of all genes in each module that mapped to significant GO pathways (padj < 0.05).

### 7. CUT&Tag-Direct

CUT&Tag-Direct was conducted with minor modifications [43, 71] [71](See CUTAC-V4 at protocols.io).

#### 7.1 Microglial nuclear isolation and preservation

Isolated microglia from two biological samples (n=2) were nucleated and lightly-cross-linked before slow-freezing. Briefly, 70 µL of isolated microglia were resuspended in 1 mL NE1 buffer (1 mL 1M HEPES-KOH pH 8.0 [ThermoFisher, J63578.AP], 500 µL 1M KCl, 12.5 µL 2M spermidine [Sigma Aldrich, 102597490], 500 µL 10% Triton X-100 [Sigma Aldrich, 1003407653], 1 Roche c0mplete Protease Inhibitor EDTA-free [Sigma Aldrich, 04693232001], and 10 mL glycerol [Sigma Aldrich, G5516-500ML] and 38 mL ddH2O) for 10 minutes on ice with light vortexing. Nuclei were centrifuged (4 minutes at 1300 *x g* at 4°C) and supernatant was removed. Isolated nuclei were then lightly crosslinked in 0.1% formaldehyde (5 mL PBS with 31 µL 16% formaldehyde (*w/v*) [ThermoFisher, 28908]) at room temperature for 2 minutes. 300 uL 1.25 M glycine [Thomas Scientific, 762Q84] was used to stop cross-linking and nuclei were centrifuged (4 minutes at 1300 *x g* at 4°C) and resuspended in 100 µL Wash Buffer (1 mL 1M HEPES pH 7.5 [ThermoFisher, J60712.AP], 1.5 mL 5M NaCl, 12.5 µL 2M spermidine, 1 Roche c0mplete Protease Inhibitor EDTA-free tablet with ddH2O to 50 mL). 10 µL of solution was taken for counting on the EVE Countess Automated Cell Counter (NanoEntek). After addition of 810 µL of Wash Buffer to resuspend nuclei, 100 µL of DMSO [ThermoFisher, 414885000] was added, followed by a brief vortex. Cross-linked microglial nuclei were then placed in a Mr. Frosty (ThermoFisher, 5100-0001) container filled with 100% isopropyl alcohol and placed at −80^°^C until future use.

#### 7.2 Concanavalin A Conjugation

Magnetic microbeads were conjugated with Concanavalin A (Sigma Aldrich, C2272-10mg) as previously described [72]. Briefly, 50 µL MyOne T1 Streptavidin-conjugated Dynabeads (ThermoFisher, 65601) were placed into a 1.5 mL tube and then placed onto a magnetic stand. The supernatant was removed, and beads were washed three times with 1 X PBS pH 6.8 (Sigma Aldrich, 1218-75) while still on the stand. Beads were then resuspended in 50 µL 1 X PBS pH 6.8 with 0.01% Tween-20 (Sigma Aldrich, 1003018242). 25 µL of biotin-conjugated Concanavalin A solution (2.3 mg/mL resuspended in 1X PBS pH 6.8 with 0.01% Tween-20) was added to the beads. Beads were incubated at room temperature for 30 minutes with rotation. Beads were placed back onto the magnetic stand and the supernatant was removed. 50 µL of 1X PBS with 0.01% Tween-20 was added to resuspend the beads.

#### 7.3 Bead Activation

Freshly conjugated beads (3.5 µL per reaction) were immediately added to 1 mL Binding Buffer (200 µL 1M HEPES pH 7.9, 100 µL 1M KCl, 10 µL 1M CaCl_2_, 10 µL 1M MnCl_2_ [ThermoFisher, J63150.AP] with 9.680 mL ddH2O) and mixed by pipetting. Beads were placed on a magnetic stand and supernatant was removed. Beads were then washed 2x with 300 µL Binding Buffer and mixed by pipetting. After removal of final supernatant, beads were resuspended in total reaction volume (i.e., 45.5 µL for 13 reactions) and left at room temperature until use.

#### 7.4 Cell Preparation and Bead Binding

Briefly, lightly cross-linked microglial nuclei were placed in room temperature water until completely thawed. Previously obtained nuclei counts were used to estimate the volume of microglial nuclei needed to achieve 2,500 nuclei per reaction (run in duplicate per mark of interest) and was added to individual 0.5 mL thin-walled PCR tubes. While gently vortexing, 3.5 µL of activated beads were added to each tube and incubated for 10 minutes at room temperature with rotation. Following incubation, samples were placed on a magnetic stand and supernatant was removed with repeated draws of a p20 pipette. Bead-bound microglial nuclei were resuspended in 25 µL Antibody Incubation Buffer (995 µL Wash Buffer with 5 µL 200x BSA [B9000S, New England Biolabs]).

#### 7.5 Primary Antibody Binding and Spike-In Preparation

Lysine-Methyl Panel Spike in (SNAP-CUTANA™ K-MetStat Panel, 19-1002, Epicypher) was diluted 1:10 in Antibody Incubation Buffer. Prior to antibody addition, 0.75 µL of 1:10 spike in panel was added to each reaction. Following this addition, 1 µL of Rabbit anti-H3K27me3 (9733S, CST), 1.5 µL of Rabbit anti-H3K27ac (8173S, CST), or 1.5 µL Rabbit (DA1E) mAb IgG (66362S, CST) antibody was added to each reaction. Samples were incubated for 2 hours at room temperature on a belly dancer (BenchWaver™ 3D Rocker, Benchmark Scientific) (105 RPM) and then transferred to 4°C for overnight incubation without rotation.

#### 7.6 Secondary and pAG-TN5 Incubation

Samples were removed from 4°C for at least 30 minutes prior to use and then resuspended using light vortexing before placing onto magnetic stand. Supernatant was removed and 25 µL of secondary antibody solution was added to each reaction (Guinea Pig Anti-Rabbit: 1:100 in Wash Buffer, ABIN101961, Antibodies Online) while gently vortexing, then incubated for 1 hour at room temperature on a belly dancer (105 RPM). Samples were then placed on a magnetic stand, supernatant removed and 200 µL Wash Buffer added without disturbing the beads. 185 µL Wash Buffer was removed, and samples spun down, followed by final liquid removal using a p20 pipette. 25 µL of pAG-TN5 adaptor complex (1:20, CUTANA™ pAG-Tn5 for CUT&Tag, 15-1017, Epicypher) in 300-Wash Buffer (1 mL 1M HEPES pH 7.5, 3 mL 5M NaCl, 12.5 µL 2M spermidine, and 1 Roche c0mplete Protease Inhibitor EDTA-free tablet with ddH2O to 50 mL) was squirted onto beads. Gently vortexing and/or tube rocking was used to ensure all beads were in solution prior to incubation for 2 hours at room temperature on a belly dancer (105 RPM).

#### 7.7 Tagmentation

Following incubation with pAG-TN5, samples were placed back onto magnetic stand and washed with 300-Wash Buffer as described previously. While taking care to avoid beads drying out, 50 µL Tagmentation Buffer (990 µL 300-Wash Buffer with 10 µL 1M MgCl_2_ [Invitrogen, AM9530G]) was added to each tube. Tubes were lightly rocked to resuspend all beads into solution followed by a brief spin-down. Samples were then incubated for 1 hour at 37°C on a thermal cycler with a hold at 8°C.

#### 7.8 Chromatin Release, Amplification and Clean-up

Samples were allowed to warm to room temperature and were gently vortexed to resuspend beads before placing back onto magnetic stand. Tagmentation Buffer was withdrawn with subsequent draws from a p20 pipette, and samples were resuspended in 50 µL TAPS Wash Buffer (30 µL 1M TAPS pH 8.5 [BostonBio Products, BB-2375], 1.2 µL 0.5M EDTA [Invitrogen, AM9262] and filled to 3 mL with ddH2O) and gently rocked to resuspend beads. Samples were placed back on magnetic stand and buffer was removed following the previous method one sample at a time. After all liquid was removed, samples were resuspended in 5 µL SDS-Release Solution (10 µL 10% SDS [ThermoFisher, 15553027], 10 µL 1M TAPS pH 6.8 and 980 µL ddH2O). A p10 pipette tip was used to drag the solution around the sides of the tubes to grab remaining beads and samples were spun down before being incubated on a thermocycler set to 58°C for 1-hour. Samples were then removed from the thermocycler, placed on ice and spiked with 15 µL of Triton Neutralization Solution (67 µL 10% Triton X-100 with 933 µL ddH2O). Samples were vortexed on high and placed back on ice. 2 µL of an i5 (10 µM) and i7 (10 µM) index [73] were then added to each reaction along with 25 µL NEBNext HiFi 2X PCR Master Mix (New England Biolabs, M0541). Samples were again vortexed on full speed for 10 seconds and briefly spun down. PCR was conducted with the following parameters: Step 1 = 58°C for 5 minutes, Step 2 = 72°C for 5 minutes, Step 3 = 98°C for 5 minutes (Cycle time increased from 30 seconds to account for full reversal of cross-linking), Step 4 = 98°C for 10 seconds, Step 5 = 60°C for 10 seconds, (Repeat Steps 4-5 14 times), Step 6 = 72°C for 1 minute and hold at 8°C. Samples were allowed to warm to room temperature for 30 minutes and then were incubated with 65 µL SPRIselect beads (Beckman Coulter, B23318) (1.3X) for 10 minutes. Samples were then placed onto a magnetic stand and beads were washed twice with 80% freshly made EtOH without removing the tubes from the stand. Samples were spun down following the last wash and placed back onto the stand to remove all traces of EtOH. The purified DNA product was released from the beads by incubating in 17 µL freshly made 10 mM Tris-HCl pH 8.0 (EMD Millipore, 648314-100mL) for 10 minutes. Samples were then placed back onto a magnetic stand and 15 µL of purified product was transferred to barcoded tubes for storage.

#### 7.9 Sample Pooling and DNA Sequencing

Samples were pooled to equimolar concentrations based on Qubit 1x dsDNA high sensitivity (ThermoFisher, Q33230) concentration and fragment size from the Bioanalyzer HS DNA kit (Agilent, 5067-4626) and sequenced on the Illumina NovaSeq6000 platform (150×150bp PE), targeting 10 million reads per sample by the University of Miami Center John P. Hussman Institute for Human Genomics sequencing core facility.

### 8. CUT&Tag-Direct Data Processing

CUT&Tag-Direct data used in this study were mapped to the mm10 genome. Prior to mapping, raw sequencing reads were mapped to the genome using Bowtie2 [74] (v. 2.4.5) with the following parameters: --local --very-sensitive --no-mixed --no-discordant --phred33 -I 10 -X 700. Resulting sam files were sorted and duplicates were marked. For control samples, (i.e. IgG targets) duplicates were removed, while remaining samples retained all duplicates. Samtools [75] (v. 1.12) was used to assess fragment length and write uniquely mapped reads from sam files into bam files. Resulting bam files were sorted and indexed. Technical replicates were kept independent and merged using samtools merge followed by sorting and indexing. Bedtools (v 2.30.0) [76] bamtobed function was then used to convert both merged files and independent files to bed files. Resulting bed files were cleaned and fragment columns were extracted using 500bp bins across the genome. These resulting files were used to assess replicate reproducibility via correlation plots. Following visualization, cleaned bed files containing fragments were normalized to barcoded read counts generated from spike-in dNuc’s with a constant value of 500,000. Peak calling was conducted using SEACR [44] on normalized bedgraph files for each biological replicate (n=2) against merged IgG controls using the following settings: “non” and “stringent”. Resulting significant peaks were used for downstream analyses. Visualization and QC plots were generated in R. All associated code for pre-processing of data files is available upon request.

### 9 Correlating RNA-sequencing Data with CUT&Tag-Direct Peaks

#### 9.1 Generation of Gene Expression Lists

Fragments Per Kilobase of transcript per Million mapped reads (FPKM) generated using Stringtie (v 2.1.7) [64] during pre-processing from RNA-sequencing data was merged and annotated using biomaRt (v 2.28.0) [77, 78] with ENSMBL ID’s. Protein-coding genes were selected and filtered based on expression levels. Genes with > 10 FPKM were subset into a “High Expression” group, genes with 1 < FPKM < 10 were subset into a “Medium Expression” group and genes with < 1 FPKM were subset into the “Low Expression” group based on brain region.

#### 9.2 Generation of Significant Called Peaks Lists

Significantly called peaks for each brain region and histone mark, based on 2 technical replicates, were annotated using HOMER [79] against the mm10 genome and any peaks mapped to genes that fall within the promoter, promoter-transcriptional-start-site (TSS) or TSS and that were protein-coding were retained for downstream analysis for H3K27ac. The same was conducted for H3K27me3 peaks, however peaks falling into an exon of these genes were also included. Fragment counts from associated bam files were used to count fragments in significantly called peaks to generate peak expression matrices for use with DESeq2.

#### 9.3 Gene Overlap

The R package GeneOverlap (v1.30.0) [80] was then used to conduct statistical overlap testing of highly expressed genes, medium expressed genes and lowly expressed genes with the presence of significant peak calls in such regions for both H3K27ac and H3K27me3. Odds ratios were used to generate intersection lists of significantly associated peaks with gene expression for further analysis. Peaks found to overlap with gene expression in both replicates were retained. To determine the epigenetic identities specific to each region, peaks found to overlap with expression in the same genes in more than one region were removed, leaving behind only those specific to one region of the brain. GO analysis was then conducted on these genes using methods described above to determine the distinct features of epigenetic regulation in each region and the rrvgo package [68] was utilized to reduce GO terms to parent terms for clarity. To generate significant overlaps between significant gene expression and significantly correlated epigenetic features, the lists of significantly differentially expressed genes with a positive or negative log_2_FC from DESeq2 conducted on RNA-sequencing data sets was taken and compared to peaks meeting the criteria as described above for highly expressed genes and presence of H3K27ac peaks, and downregulated genes and the presence of H3K27me3 peaks. The number of genes intersecting were used to generate representative Venn diagrams. All protein coding genes mapped were used as background for gene overlap.

#### 9.4 Visualization of Overlapping CUT&Tag-Direct/RNA-sequencing data

Bam files generated from mapped read alignments for both RNA-sequencing and CUT&Tag-Direct sequencing data were used to generate bigWig files for visualization. Samples 7 and 8 from RNA-sequencing data were used as input as they correspond to the same animals and regions used for CUT&Tag-Direct data. Using deepTools [81] bamCoverage, bam files from CUT&Tag-Direct sequencing were screened against blacklisted regions [82] and converted to bigWig files using the following parameters: --binSize 10, -- normalizeUsing RPCG – effectiveGenomeSize 2652783500 –ignoreForNormalization chrX and -- extendReads. Following conversion, bigWig files from biological replicates were merged using bigWigMerge from the UCSC genome browser bwtools. Merged files were sorted and then converted back into bigWigs using the bedgraphToBigWig program [83]. For RNA-sequencing data, using deepTools [81] bamCoverage, bam files were normalized for read coverage using the inverse scale factors calculated by DESeq2 and converted to bigWig files using the following parameters: --binSize 1 and --skipNonCoveredRegions. Following conversion, bigWig files from biological replicates were merged using bigWigMerge from bwtools. Merged files were sorted and then converted back into bigWigs using the bedgraphToBigWig program [83]. The resulting bigWig files were visualized using rtracklayer (v 1.58.0) [84], TrackPlotR (v. 0.0.1) [85] and ggplot2 (v. 3.5.0.9000).

### 9. Statistical Analysis

Statistical analyses were conducted in accordance with the various packages used for RNA-sequencing data as well as CUT&Tag-Direct data. Information on specific analyses run can be found in the associated code and methods sections.

## Funding

DP1DA051828 (LMT)

K01DA045294 (LMT)

Kind gift from the Shipley Foundation (LMT)

## Author contributions

Conceptualization: AVM, SJV, LMT

Methodology: AVM, SJV, FBG, LMT

Software: AVM

Investigation: AVM, SJV, LMT

Visualization: AVM, SJV

Data Curation: AVM, SJV

Supervision: FBG, LMT

Writing—original draft: AVM, SJV, LMT

Writing—review & editing: AVM, SJV, LMT

## Data and materials availability

All next generation sequencing files associated with this study as well as the code that was used to pre-process and run differential expression are available online at https://github.com/avm27/RegionalMicroglialTranscriptomicsandEpigenomics. All data files used in processing and analysis will be available from the GEO/SRA following publication and/or upon request.

