## Supplementary Materials for "Epigenetic heterogeneity shapes the transcriptional landscape of regional microglia"

Alexander V. Margetts *et al.*

**This PDF file includes:**

Figs. S1 to S3
Data S1 to S3

Fig. S1.

A

**Reads on-target for H3K27me3**

| Target | Negative Control | PFC8 | PFC9 | STR8 | STR9 | HPP8 | HPP9 | VTA8 | VTA9 | CTX8 | CTX9 |
|---|---|---|---|---|---|---|---|---|---|---|---|
| Unmodified | 5.68% | 0.84% | 2.85% | 1.04% | 0.98% | 1.05% | 1.42% | 0.64% | 0.71% | 1.71% | 2.02% |
| H3K4me1 | 6.99% | 1.13% | 4.68% | 0.88% | 1.09% | 0.97% | 1.46% | 0.67% | 0.83% | 2.42% | 1.53% |
| H3K4me2 | 6.85% | 1.25% | 3.28% | 1.20% | 1.01% | 1.26% | 1.62% | 0.85% | 0.98% | 2.31% | 2.08% |
| H3K4me3 | 5.11% | 1.73% | 3.70% | 1.65% | 2.25% | 3.23% | 3.20% | 1.58% | 1.74% | 4.02% | 2.87% |
| H3K9me1 | 6.20% | 1.10% | 3.27% | 0.94% | 1.03% | 1.12% | 1.35% | 0.88% | 0.90% | 2.66% | 1.82% |
| H3K9me2 | 4.37% | 0.80% | 4.23% | 0.69% | 0.85% | 0.73% | 1.04% | 0.52% | 0.65% | 1.74% | 1.70% |
| H3K9me3 | 4.96% | 3.26% | 7.11% | 3.19% | 5.12% | 8.93% | 5.66% | 3.95% | 4.35% | 8.35% | 6.13% |
| H3K27me1 | 4.19% | 0.60% | 3.61% | 0.69% | 0.70% | 0.58% | 1.48% | 0.37% | 0.61% | 1.81% | 1.20% |
| H3K27me2 | 4.71% | 2.17% | 3.74% | 1.88% | 2.52% | 2.72% | 3.49% | 1.57% | 1.86% | 4.82% | 3.55% |
| H3K27me3 | 5.78% | 100.00% | 100.00% | 100.00% | 100.00% | 100.00% | 100.00% | 100.00% | 100.00% | 100.00% | 100.00% |
| H3K36me1 | 7.56% | 1.03% | 2.84% | 1.04% | 1.09% | 1.11% | 2.13% | 0.79% | 0.76% | 2.27% | 2.34% |
| H3K36me2 | 8.99% | 1.76% | 4.77% | 1.32% | 1.40% | 1.45% | 2.42% | 1.03% | 1.46% | 2.70% | 2.73% |
| H3K36me3 | 7.87% | 1.70% | 3.70% | 1.51% | 1.60% | 1.66% | 2.77% | 1.20% | 1.19% | 3.16% | 1.87% |
| H4K20me1 | 8.89% | 1.56% | 3.26% | 1.43% | 1.51% | 1.38% | 2.18% | 1.03% | 0.96% | 2.77% | 1.74% |
| H4K20me2 | 8.03% | 1.20% | 3.09% | 1.21% | 1.28% | 1.14% | 2.35% | 0.88% | 1.13% | 2.38% | 2.13% |
| H4K20me3 | 3.82% | 0.65% | 2.02% | 0.53% | 0.63% | 0.71% | 1.41% | 0.43% | 0.46% | 1.58% | 1.70% |
| % total reads | 7.794% | 0.106% | 0.079% | 0.052% | 0.064% | 0.134% | 0.075% | 0.081% | 0.142% | 0.086% | 0.061% |

B

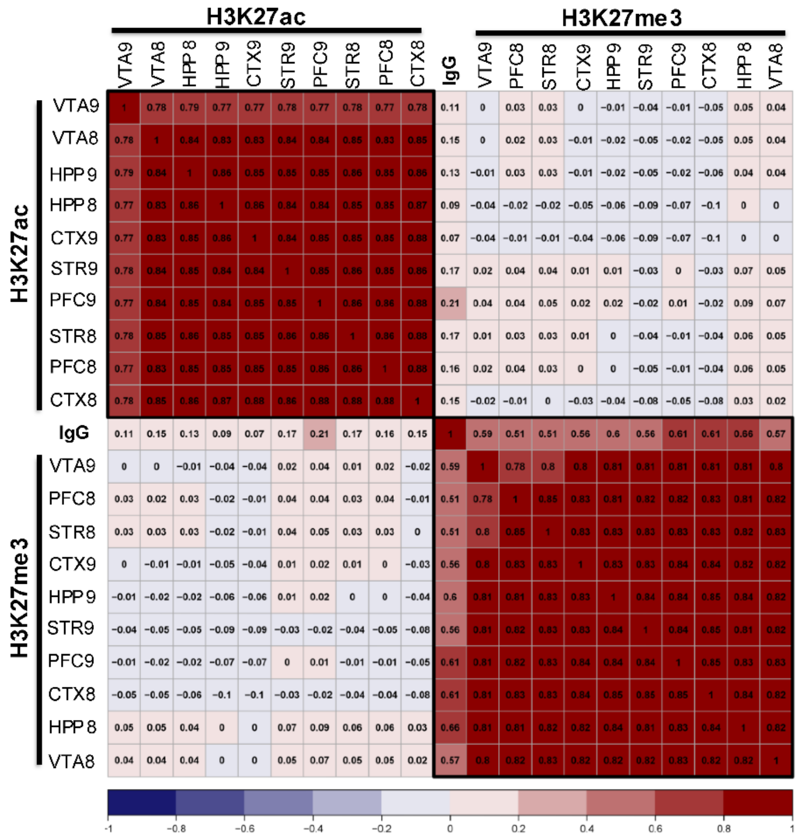


**CUT&Tag-Direct is specific and reproducible. (A)** Heatmap showing the normalized percentage of reads mapped to dNuc's (Epicypher) for H3K27me3 with colors indicating the relative rates of mapping. **(B)** Correlation plots indicating the relative similarity of uniquely mapped 500bp fragments for H3K27ac, H3K27me3 and IgG.

**Fig. S2.**

A

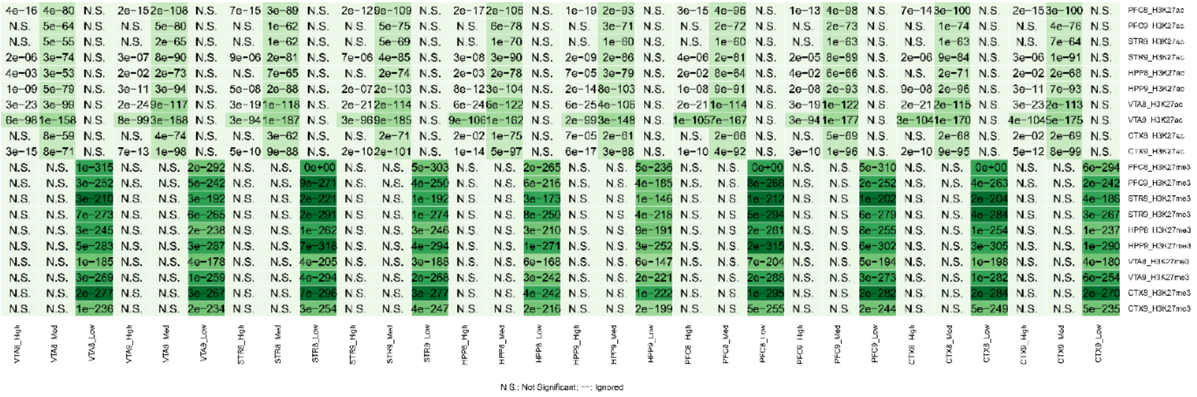


B

| Region | Histone Mark | Overlapping Genes | Overlapping Unique Genes | Percent Uniqueness |
|---|---|---|---|---|
| PFC | H3K27me3 | 1960 | 229 | **11%** |
| STR | | 1190 | 146 | **12%** |
| VTA | | 1152 | 213 | **18%** |
| HPP | | 1609 | 186 | **11%** |
| CTX | | 687 | 194 | **28%** |
| PFC | H3K27ac | 738 | 151 | **20%** |
| STR | | 667 | 117 | **17%** |
| VTA | | 897 | 267 | **29%** |
| HPP | | 739 | 173 | **23%** |
| CTX | | 749 | 162 | **21%** |

**Gene overlap analysis of significant peaks in differentially expressed genes. (A)** Odds ratio of significantly called peaks for each mark (H3K27ac or H3K27me3) and region against gene expression (Low: FPKM < 1, Medium: FPKM > 1 and < 10, High: FPKM > 10, padj shown). **(B)** Table showing the number of significantly overlapping genes with either low expression (FPKM < 1) and called peaks for H3K27me3 or high expression (FPKM > 10) and called peaks for H3K27ac. Percent uniqueness was calculated by selecting only those genes that are either lowly expressed and have deposition of H3K27me3 or highly expressed and have deposition of H3K27ac and are found in only one region divided by all overlapping genes in that region.

**Fig. S3.**

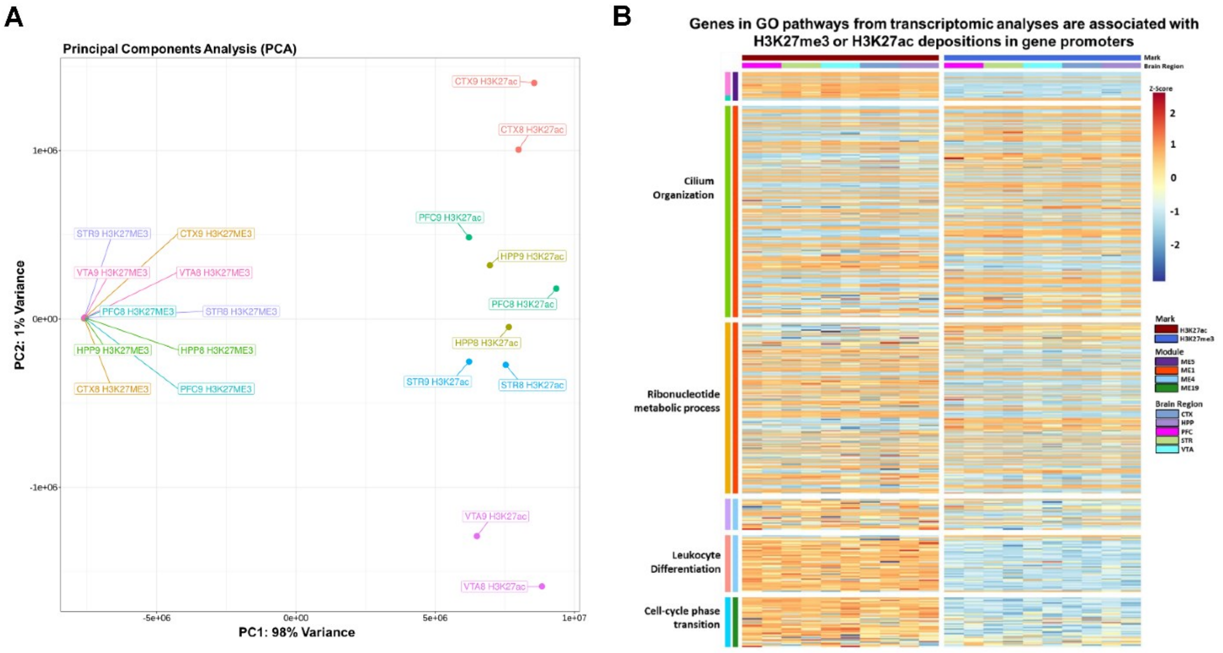


**H3K27acetylation could be a driver of variation in regional microglia. (A)** PCA of fragments of peaks in genes for either H3K27me3 or H3K27ac. **(B)** Heatmap of genes identified in WGCNA modules and corresponding enriched go pathways (padj < 0.05), showing the levels of H3K27me3 or H3K27ac in promoters or exons of such genes.

**Data S1. (separate file)**

**SupplementaryDataFile1.xlsx**

Top 50 up and down regulated genes in transcriptional analysis based on comparison after differential gene expression analysis.

**Data S2. (separate file)**

**SupplementaryDataFile2.xlsx**

Results for WGCNA analysis including statistical values, module expression and statistically significant GO pathways.

**Data S3. (separate file)**

**SupplementaryDataFile3.xlsx**

Alignment and QC statistics for CAT data and peak calling parameters. Significant peak annotations for H3K27me3 and H3K27ac peak calling files and all deseq2 results including top 50 up and down differential depositions of peaks and all identified GO pathways.
